# *iucn_sim*: A new program to simulate future extinctions based on IUCN threat status

**DOI:** 10.1101/2019.12.16.878249

**Authors:** Tobias Andermann, Søren Faurby, Robert Cooke, Daniele Silvestro, Alexandre Antonelli

## Abstract

The ongoing environmental crisis poses an urgent need to forecast the *who*, *where*, and *when* of future species extinctions, as such information is crucial for targeting conservation efforts. Commonly, such forecasts are made based on conservation status assessments produced by the International Union for Conservation of Nature (IUCN). However, when researchers apply these IUCN conservation status data for predicting future extinctions, important information is often omitted, which can impact the accuracy of these predictions.

Here we present a new approach and a software for simulating future extinctions based on IUCN conservation status information, which incorporates generation length information of individual species when modeling extinction risks. Additionally, we explicitly model future changes in conservation status for each species, based on status transition rates that we estimate from the IUCN assessment history of the last decades. Finally, we apply a Markov chain Monte Carlo algorithm to estimate extinction rates for each species, based on the simulated future extinctions. These estimates inherently incorporate the chances of conservation status changes and the generation length for each given species and are specific to the simulated time frame.

We demonstrate the utility of our approach by estimating extinction rates for all bird species. Our average extinction risk estimate for the next 100 years across all birds is 6.98 × 10^−4^ extinctions per species-year, and we predict an expected biodiversity loss of between 669 to 738 bird species within that time frame. Further, the rate estimates between species sharing the same IUCN status show larger variation than the rates estimated with alternative approaches, which reflects expected differences in extinction risk among taxa of the same conservation status. Our method demonstrates the utility of applying species-specific information to the estimation of extinction rates, rather than assuming equal extinction risks for species assigned to the same conservation status.

## Introduction

We are in the middle of a massive biodiversity crisis (Barnosky et al. 2011, Davis et al. 2018, Díaz et al. 2019). Extinction risks have been steadily increasing for as long as we have been keeping record, with no indications of a slowdown (Ceballos et al. 2015). It is therefore crucial to predict the number of future extinctions that shape the future biodiversity, whether in terms of species, phylogenetic, or functional diversity (Davis et al. 2018, Cooke et al. 2019, Pimiento et al. 2020). An important use of such predictions is to aid conservation prioritization (Mooers et al. 2008). However, all predictions require reliable estimates of extinction risk.

The main global initiative to quantify extinction risks across animal and plant species is the IUCN Red List (IUCN Red List 2019), which categorizes the conservation status of organisms based on expert assessments. Since 2001, the IUCN has adopted the IUCN v3.1 evaluation system for determining species’ conservation statuses (IUCN Species Survival Commission 2001). By this standard, extant species are assessed as Least Concern (LC), Near Threatened (NT), Vulnerable (VU), Endangered (EN), or Critically Endangered (CR). If there is insufficient information available for a species to enable a proper status assessment, the species is categorized as Data Deficient (DD). Species that have not yet been reviewed by the IUCN are categorized as Non-Evaluated (NE). Species that are not found in the wild anymore are labeled as Extinct in the Wild (EW), and species with no living wild or captive individuals as Extinct (EX). As of the year 2020, IUCN has also introduced two additional subcategories for CR species (IUCN 2020): possibly extinct [CR(PE)], and possibly extinct in the wild [CR(PEW)].

For the IUCN to decide on assigning a species to one of the threatened categories VU, EN, or CR, this species must meet at least one of five assessment criteria (A-E). One of those criteria (E) is associated with a specific extinction probability, while the other criteria (A-D) mostly encompass estimates of decreasing population trends and fragmentation. The IUCN extinction probability thresholds defined in criterion E are as follows:

- VU: 10% extinction probability within 100 years
- EN: 20% extinction probability within 20 years or 5 generations, whichever is longer (maximum 100 years)
- CR: 50% extinction probability within 10 years or 3 generations, whichever is longer (maximum 100 years)

IUCN conservation status assessments have been used in numerous scientific studies to project future biodiversity loss (e.g. Ricciardi and Rasmussen 1999, Veron et al. 2016, Davis et al. 2018, Cooke et al. 2019, Oliveira et al. 2019). One critical challenge in this approach is to meaningfully transform the IUCN-defined conservation statuses into explicit extinction probabilities. In these previous studies, researchers have applied the extinction probabilities associated with criterion E to model extinction risks for threatened species. Sometimes these risks are also extrapolated to species of the statuses LC and NT in order to make it possible to assign extinction probabilities to these species (Redding and Mooers 2006, Mooers et al. 2008, Veron et al. 2016, Davis et al. 2018).

Although these extinction probabilities only apply to species that are assessed under criterion E (see Akçakaya et al. 2006), they are commonly applied equally to all species sharing the same conservation status (e.g. Mooers et al. 2008, Davis et al. 2018). The underlying assumption that the minimum extinction risks defined for criterion E can be meaningfully transferred to species listed under one of the other four criteria (A-D) is difficult to test empirically, but is a necessary simplification in order to model the extinction probabilities for the majority of species. However, there are several other important aspects that can be easily incorporated but are commonly neglected when translating IUCN conservation statuses into extinction probabilities for future extinction predictions.

### Neglected information

To the best of our knowledge, there are two key elements that are usually not incorporated when using IUCN status data for future extinction predictions: generation length and expected changes in conservation status (but see Monroe et al. 2019).

Generation length (GL) is defined as the average turnover rate of breeding individuals in a population (IUCN Standards and Petitions Committee 2019) and therefore reflects the turnover between generations. It is generally considered to be a more meaningful time unit for modeling extinction risk than time expressed in years (Frankham and Brook 2004). Generation length should not be confused with age of sexual maturity, which can be used in the calculation of generation length, but is not equivalent (with age of sexual maturity always being smaller than or equal to generation length). As per the IUCN definition, the extinction probability for the categories EN and CR is to be understood in context of the GL of the given species, if 5 × *GL* exceeds 20 years for EN species, or if 3 × *GL* exceeds 10 years for CR species (see criterion E definitions above). We argue that including GL should be the standard practice when modeling extinction risks based on IUCN data, particularly because GL data is readily available for many species (Pacifici et al. 2013, BirdLife International 2019, IUCN Red List 2019) and can normally be approximated through a combination of body size and phylogenetic information (Cooke et al. 2018, Bird et al. 2020) for species missing GL data (Appendix 1).

A second missing element in many future predictions relates to the fact that IUCN categories are generally treated as static entities that do not change over time. However, almost two decades of IUCN re-assessments of species (IUCN Red List 2019), using the IUCN v3.1 standard, have clearly shown this not to be the case. Instead, re-assessments (Butchart et al. 2007, Rondinini et al. 2014) show that the conservation status of species can change significantly in a relatively short time span, for instance as a result of the effectiveness of conservation efforts. For a species classified as LC, the immediate extinction risk is negligibly small, while for a species classified as CR, the immediate extinction risk is very high. It is reasonable to assume that if we simulate, for example, 100 years into the future, categories may change due to new or intensified risks or thanks to conservation efforts, which will affect the extinction probabilities.

An example of a change in IUCN status is the Mauritian Pink Pigeon (*Nesoenas mayeri*), which was listed as CR in the 1990s, with only 9 birds remaining, due to habitat loss and predation by introduced species (Swinnerton 2001, IUCN Red List 2019). However, following an intensive conservation recovery program, the Pink Pigeon is now listed as VU, with around 470 wild birds (IUCN Red List 2019). Unfortunately, most species show changes with the opposite trend, for example several species of vultures, which are declining due to poisoning and persecution (Green et al. 2007). There are 22 species of vulture (Accipitridae: Gypaetinae, Accipitridae: Aegypiinae, and Cathartidae) according to the IUCN Red List, 12 of these are classified as threatened (VU, EN or CR), including 9 CR species (IUCN Red List 2019), with sharp declines in population sizes. For instance, four species of vultures (the White-headed Vulture *Trigonoceps occipitalis*, White-backed Vulture *Gyps africanus*, Hooded Vulture *Necrosyrtes monachus* and Rüppell’s Vulture *Gyps rueppelli*) were all listed as LC in 2004 but are now all classified as CR. Information about these changes can be accessed through the IUCN history record and can then be used to inform models of extinction risk.

While many previous studies have applied the IUCN-based extinction probabilities (criterion E) outlined above to model extinction risks, a recent study by Monroe et al. (2019) has presented an alternative approach, avoiding these probabilities altogether. Instead Monroe et al. (2019) modeled extinction risks based on observed transitions of species to the statuses EW or EX, which they then applied to model future biodiversity losses. While this approach avoids the above mentioned shortcomings of the IUCN extinction probability approach, namely the caveats surrounding the extrapolation of the criterion E specific extinction probabilities to species listed under other criteria (A-D), it is likely limited to groups of organisms with sufficient recorded transitions to EX in order to yield reliable extinction risk estimates.

In this study we contrast different variations of both approaches, to which we refer to hereafter as “critE EX mode” (approach based on IUCN criterion E extinction probabilities, sensu Mooers et al. 2008) and “empirical EX mode” (approach based on historic transitions towards statuses EW/EX, sensu Monroe et al. 2019). We add improvements to both approaches, including the incorporation of GL data for the critE EX mode and the consideration of possibly extinct taxa in the empirical EX mode. Further we present a novel MCMC-based approach of estimating status transition rates from historical IUCN data, and we apply these rates to simulate future status changes and extinctions. All approaches presented in this study are available in the new open-access simulation program ***iucn_sim***, which can be run via the bash command line and is tested for compatibility in Windows, MacOS, and Linux (Fig. 1, see Data availability statement).

**Figure 1:**
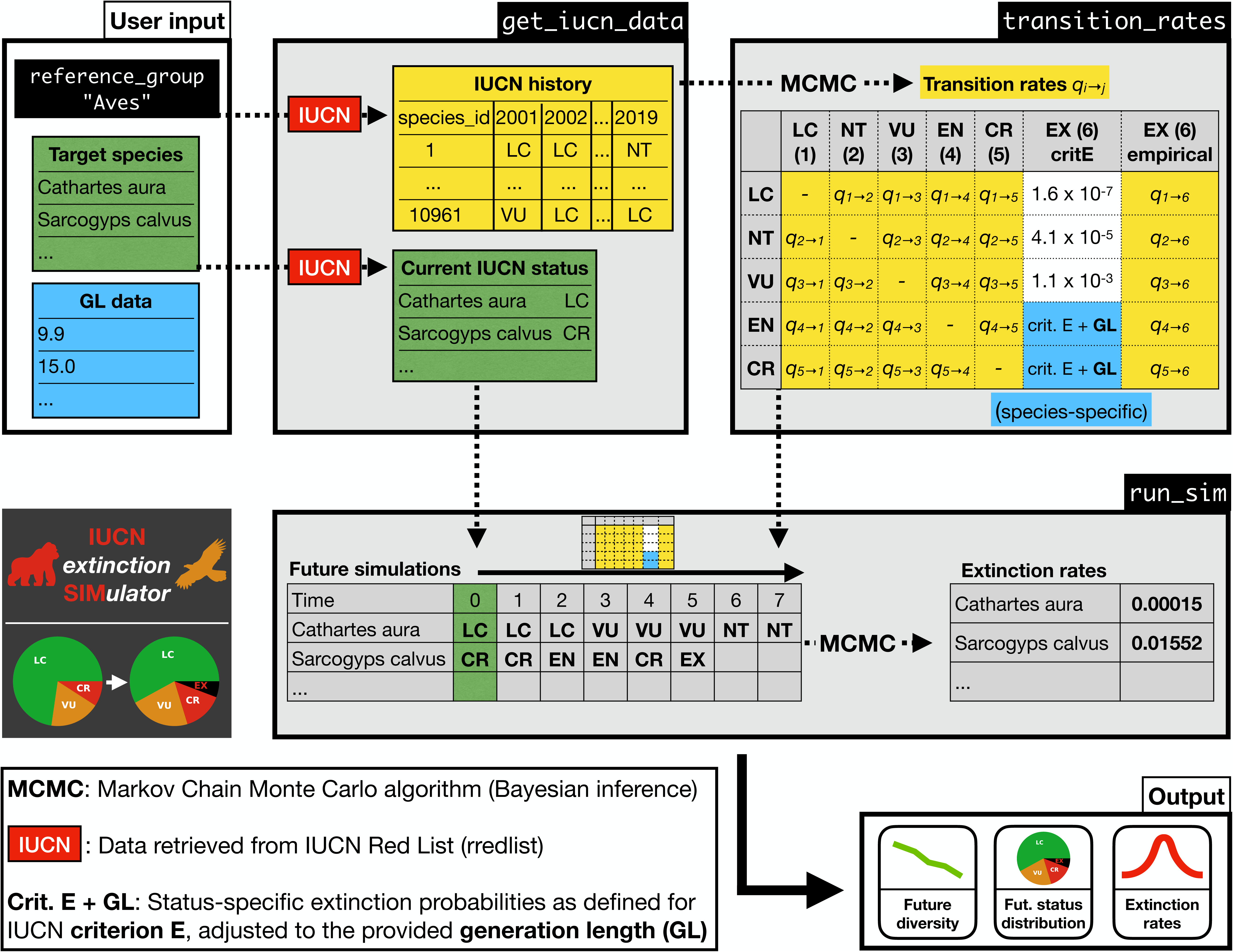
Workflow of ***iucn_sim*** to simulate future extinctions and estimate extinction rates. The only required input by the user is a) the list of target species whose future extinctions are supposed to be simulated and b) the name of a reference group, which will be used to estimate status transition rates based on the recorded IUCN history of this group. Optionally the user can provide generation length (GL) estimates for each target species, which will be considered when calculating the extinction risks associated with the statuses EN and CR, according to IUCN criterion E (critE EX mode). Alternatively, the user can choose the empirical EX mode, in which case extinction risks will be estimated from the empirically observed extinctions in the IUCN history of the reference group. The modeled extinction risks and the status transition rates will be stored in a q-matrix, which is used to simulate future status changes and extinctions for all target species. Finally, the program estimates species-specific extinction rates from the simulated extinction times (typically from multiple simulation replicates) and produces various summary statistics and plots as output, including the simulated future status distribution of the target group, the future diversity trajectory, and the histograms of simulated extinction times and extinction rate estimates for each species.

## Material and methods

Here we describe our approach of simulating future extinctions and IUCN status transitions on the example of birds (Aves). We use the terms “reference group” and “target species” as follows:

- **Reference group**: The group of species, whose IUCN history is being used to estimate status transition rates, i.e. the rates at which species change between IUCN statuses
- **Target species**: The group of species, for which future extinctions are being simulated, while applying the estimated status transition rates

In this study the reference group and target species list consist of the same taxa (all extant bird species), but this is not a requirement for this approach. For example, using our approach one could simulate future extinctions and status transitions for a specific bird family or local bird fauna, while using all birds as a larger reference group, in order to get reliable status transition rate estimates.

To make this approach accessible and easy to use for future projects, we wrapped the complete workflow described below into the open-source command line program ***iucn_sim*** (Fig. 1, https://github.com/tobiashofmann88/iucn_extinction_simulator), which can be easily installed together with all software dependencies using the conda package manager (https://docs.conda.io). Installation instructions are available on the projects GitHub page of this project. Using ***iucn_sim***, it is straight-forward to 1) model future changes in IUCN status, 2) simulate possible times of extinction across species, and 3) estimate species-specific extinction rates for any given set of species over a user-defined time span. We executed all steps outlined below using ***iucn_sim*** (except the downloading of GL data) and we report the ***iucn_sim*** command for each step in the Supplementary Code Sample 1.

### Generation length estimates

We downloaded GL estimates for all bird species from Bird et al. (2020), following the IUCN 2019-v2 taxonomy of extant bird species. Since the collecting of GL data can be challenging for some groups, we provide instructions how to generate GL data via phylogenetic imputation on the example of birds in Appendix 1 in the Supplementary Material. See Cooke et al. (2018) and Bird et al. (2020) for more detailed instructions and information on generating GL estimates for species.

### Downloading IUCN data

We downloaded the complete IUCN history for the reference group (class Aves) from the year 2001 onward, to ensure compatibility with the IUCN v3.1 standard (IUCN Species Survival Commission 2001), using the ***rl_history*** function of the R-package rredlist (Chamberlain 2017) and IUCN v2019-2. The taxon list for this download was generated by scanning through the entire IUCN Red List catalogue for species assigned to the class Aves, using the ***rl_sp*** function. In addition to the historic data, we extracted the current status (most recent status assessment) of all target species (all Aves species) using the ***rl_search*** function. For all following operations we set the status of all EW species to EX.

### Status transition rates

Based on the IUCN history data, we counted all types of status changes that have occurred in the IUCN history of birds (Table 1), as well as the cumulative amount of time spent in each status across all bird species. For instance, if a given species was classified as NT from 2001 to 2005 and then EN from 2005 to today (2020), this species contributes 1 status change from NT to EN and 4 years in NT and 16 years in EN.

**Table 1:** Status transitions counted in the IUCN history of birds (class Aves) between 2011-2020. For example, the empirical count of transitions from status LC to NT is 182, while the count of transitions from NT to LC is 112. The count for transitions from CR to EX changes from 6 to 20 when modeling species that are possibly extinct according to IUCN (PEX) as EX.

|  | LC | NT | VU | EN | CR | DD | EW/EX |
| --- | --- | --- | --- | --- | --- | --- | --- |
| LC | 0 | 182 | 77 | 18 | 3 | 1 | 0 |
| NT | 112 | 0 | 73 | 24 | 3 | 1 | 0 |
| VU | 18 | 82 | 0 | 99 | 13 | 1 | 0 |
| EN | 1 | 15 | 69 | 0 | 50 | 0 | 0 |
| CR | 0 | 2 | 9 | 42 | 0 | 0 | 6 (+14) |
| DD | 8 | 12 | 5 | 2 | 0 | 0 | 0 |
| EW/EX | 0 | 0 | 0 | 0 | 2 | 0 | 0 |

From these counts we estimated the rates of transitions between all pairs of statuses using Bayesian sampling. For example, if N_ij_ transitions were observed from status *i* to status *j* and the cumulative time spent in *i* across all species in the reference group is *t*_*i*_, we used a Markov chain Monte Carlo (MCMC) algorithm to sample the annual transition rate q_ij_ from the following posterior:

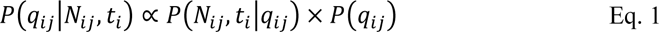

where the log likelihood function is that of a Poisson process describing status change

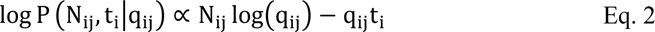

and P(q_ij_) ∼ 𝒰[0,1000] is a vague uniform prior on the transition rate.

To incorporate the uncertainties in the transition rate estimates, we took 100 samples from the posterior distribution of the rate estimates (Eq. 1) for each transition type (Table 2). More specifically we populated 100 q-matrices containing the sampled rates for each type of status transition. These q-matrices were used for future simulations, allowing us to simulate potential future status changes of any species, given its starting status (see more detailed explanation below). In addition, we sampled 100 transition rates for all transition types from DD to any of the statuses LC, NT, VU, EN, and CR, which we used during the future simulations to draw a new valid status for DD species.

**Table 2:** Status transitions rates estimated for birds (class Aves) that were used for future simulations. The q-matrix below shows the mean of the transition rate estimates across the q-matrix replicates for all bird species, scaled in transitions per species-year (T/SY). The transition rates between all extant statuses were estimated from the counted transitions in the IUCN history of birds (Table 1), taking into account the total cumulative time across all bird species spent in each category. The EX transition rates are shown for both approaches, the critE EX mode and the empirical EX mode (including PEX taxa), respectively.

|  | LC | NT | VU | EN | CR | EX |
| --- | --- | --- | --- | --- | --- | --- |
| LC | - | 0.00146420 | 0.00063157 | 0.00015742 | 0.00003312 | 0.00000016 / 0.00000794 |
| NT | 0.00755463 | - | 0.00491179 | 0.00161410 | 0.00025690 | 0.00004155 / 0.00006458 |
| VU | 0.00164326 | 0.00691254 | - | 0.00831167 | 0.00114747 | 0.00105305 / 0.00007353 |
| EN | 0.00030274 | 0.00248436 | 0.01058826 | - | 0.00757056 | 0.00900926 / 0.00014377 |
| CR | 0.00033174 | 0.00100060 | 0.00319970 | 0.01443416 | - | 0.04767378 / 0.00654700 |

Finally, we modelled the transition rates towards extinction (EX) from any extant status *i* (*q*_*i*→*EX*_). Modelling these transition rates towards EX is non-trivial and we used two approaches to estimate these rates, which we refer to as the critE EX mode and the empirical EX mode. The critE EX mode is based on the IUCN criterion E extinction probabilities defined for threatened statuses (sensu Mooers et al. 2008), whereas the empirical EX mode is based on empirically observed transitions towards EW/EX in the IUCN history (sensu Monroe et al. 2019).

The final q-matrices contain all status transition rates, including the rates towards extinction for each given status, determined with either the critE EX mode or the empirical EX mode outlined below (last column). Since in our simulations, species are not allowed to reappear after extinction, we set the rates from EX to any other status equal to 0:

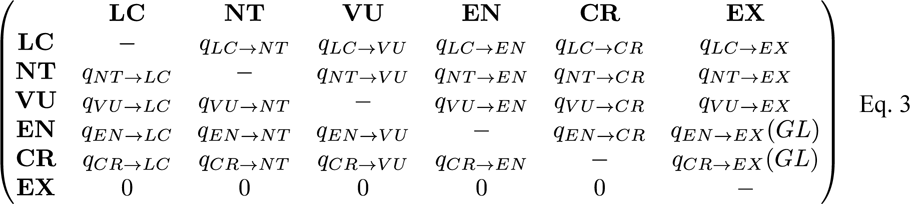

#### 1) CritE EX mode

In the critE EX mode approach (***iucn_sim*** setting: *--extinction_probs_mode 0*) we transformed the extinction probabilities (*E*_*t*_) associated with threatened IUCN statuses (see Introduction), defined over specific time frames (*t*), into annual status-specific EX transition rates (*q*_*i*→*EX*_), using the formula provided by (Kindvall and Gärdenfors 2003):

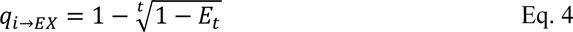

Since the IUCN extinction probabilities are only defined for the statuses VU, EN, and CR, we extrapolated the annual EX transition rates for the remaining statuses LC and NT by fitting a power function to the calculated extinction rates for the statuses VU, EN, and CR, estimating the parameters *a* and *b* (Appendix 1):

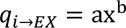

where *x* represents the index of the IUCN category, sorted by increasing threat (i.e. x_LC_ = 1, x_NT_ = 2, …, x_CR_ = 5). After estimating the parameters a and b, we calculated the annual transition rates to EX for statuses LC and NT, using the above power function.

According to the IUCN definition, the extinction probabilities linked to the IUCN categories EN and CR for individual species are dependent on the GL of these species. In order to properly model the EX transition rates for these statuses on a species-specific basis, we applied GL data to adjust the time frame (*t*) associated with the extinction probabilities. For example for a species with a GL of 5 years, which is categorized as CR (IUCN definition: “50% extinction probability within 10 years or 3 generations, whichever is longer”), the annual EX transition rate according to Eq. 4 is 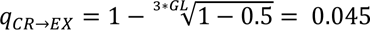, whereas for a species with a GL of 2 years with status CR the EX transition rate is *q*_*CR*→*EX*_ = 1 – 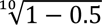 = 0.067, because in the latter case 3 ∗ *GL* < 10 *years*. From this follows that when ignoring GL information and setting *t* = 10 for all species (e.g. Mooers et al. 2008), the extinction risk for species with moderate or long generation times (>3.33 years) will be overestimated (Fig 2).

**Figure 2:**
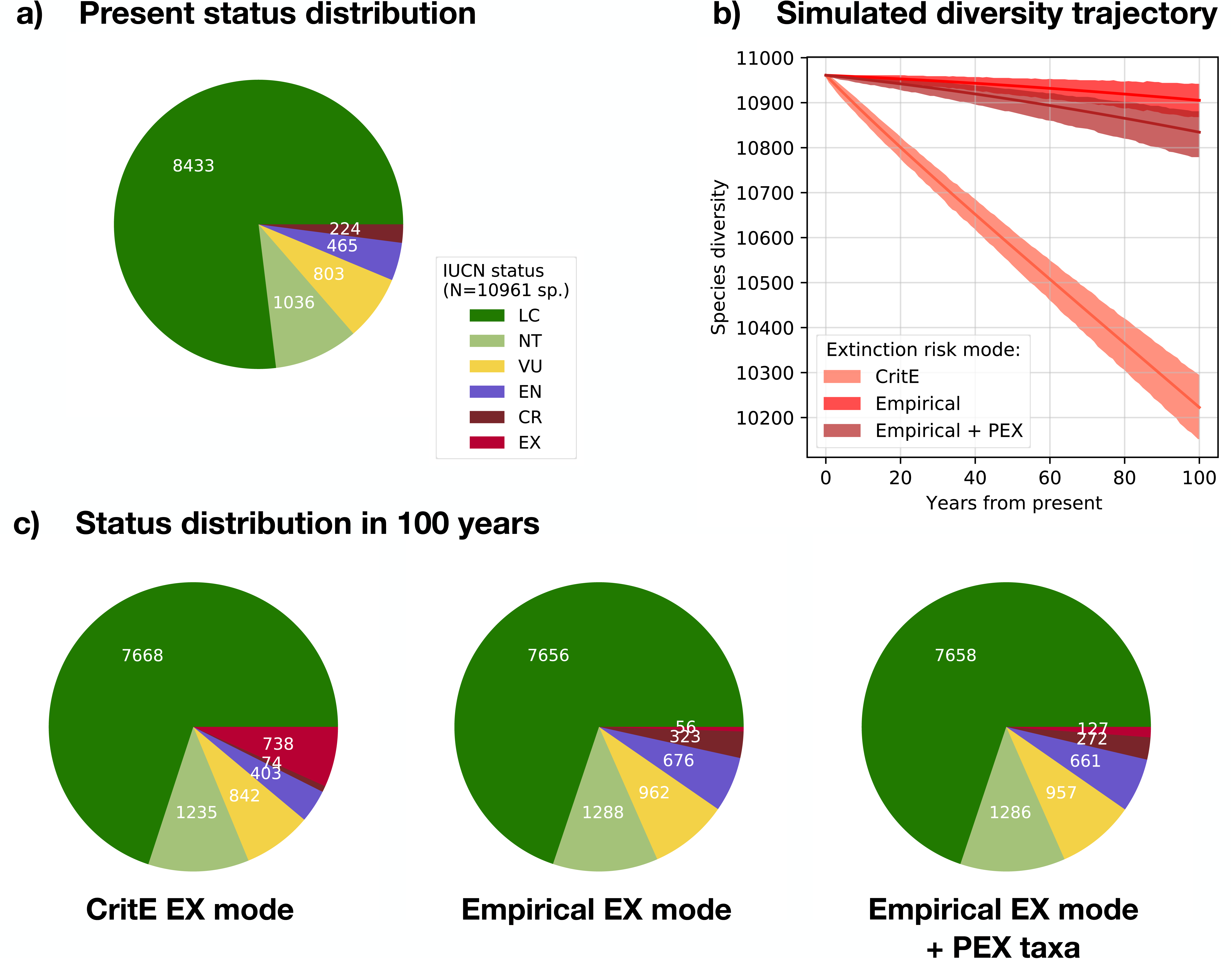
Future diversity trajectory and IUCN status distribution for birds. We simulated future extinctions with three different approaches of modeling extinction risks: the critE EX mode, the empirical EX mode, and the empirical EX mode including the modeling of PEX species as extinct. Panel a) shows the current IUCN threat status distribution of all bird species. Panel b) shows the future diversity trajectory over the next 100 years, based on future extinctions simulated with ***iucn_sim*** under the 3 different extinction risk scenarios. Panel (c) shows the future IUCN status distribution in 100 years simulated with ***iucn_sim***. Note that the y-axis in the diversity through time plots only displays a selected diversity range starting at 10,000 (displayed range does not cover the value 0). All panels represent graphic output options available in ***iucn_sim***.

We therefore applied the GL estimates of all individual bird species (Bird et al. 2020) for the calculation of the yearly EX transition rates for the statuses EN and CR of each species. We then added these GL-adjusted EX transition rates to the q-matrices containing the extant status transition rates (Eq. 3) to simulate future status changes and extinctions. Although the EX transition rates derived in this manner and the extant status transition rates are modelled based on two different data sources, they can be combined in the same q-matrix since all of these rates are expressed in the same unit (transition events per species and year). To evaluate how the presence of GL data effects future extinction predictions, we produced an additional set of q-matrices where we did not apply GL data (“no GL” scenario). Further, to evaluate the effect of modeling future status changes, we produced additional sets of q-matrices where all transition rates between extant statuses were set to 0 (“no status change” scenarios), for both the GL and no GL scenario.

#### 2) Empirical EX mode

In the empirical EX mode approach (***iucn_sim*** setting: *--extinction_probs_mode 1*) we estimated EX transition rates based on the observed transitions from any given extant status to EX in the IUCN history of birds, *sensu* Monroe et al. (2019). Following the same procedure we used to infer transition rates between other statuses, we counted the transitions from any status to EX (Table 1) and used MCMC to sample from the posterior transition rate distribution (Eq. 1), of which we randomly selected 100 samples for each type of transition. In contrast to the approach of Monroe et al. (2019), we estimated a specific transition rate from any given status to EX using our MCMC based approach, instead of only allowing transitions from CR to EW/EX. However, due to no observed occurrences of transitions of the statuses LC, NT, VU, and EN to EX in the IUCN history of birds, the estimated rates for these types of transitions are negligible, effectively making these transitions events very unlikely in our simulations of future extinctions. A further difference is that we did not distinguish between the statuses EW and EX, but instead treated both statuses as extinct. We inserted the 100 sampled rates into the last column of the q-matrix (Table 2). Since no GL data replicates or other species-specific data were used in this approach, the same 100 q-matrices were applied to all species for future simulations.

The empirical EX mode approach is likely to underestimate the true transition rate, as for several threatened species there is insufficient evidence to categorize them as EX, although they are likely to be extinct (IUCN 2020). In order to better account for this underestimation bias, we generated another set of q-matrices, where we incorporated information on possibly extinct species [CR(PE) *sensu* IUCN 2020]. Prior to determining the numbers of transitions, we modeled these species as EX, starting from the date that they were categorized as PEX (the empirical EX mode + PEX approach). This modeling of PEX species was only done for the purpose of estimating EX transition rates, while these species were classified as CR as a departure point for future simulations (see below).

### Simulating future extinctions

We used the transition rates from the estimated q-matrices (Eq. 3) to simulate for all bird species future transitions between extant statuses or to toward EX. Before simulating into the future, each species was assigned its current IUCN status as starting status. For all species currently assigned as DD, we randomly drew a new status based on a probability vector consisting of the estimated transition rates leading from DD to the valid statuses LC, NT, VU, EN, or CR. For all NE species we drew a new valid IUCN status based on the frequencies of valid IUCN statuses across all bird species. We treated species that are categorized as EW or EX by IUCN as irrevocably extinct and therefore did not include these species in future simulations.

We modeled transitions as a Poisson process, by generating time-forward simulations for each species based on exponentially distributed waiting times between transition events. For a given current status *i* the waiting time until the next event is

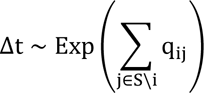

where *S*\*i* is the set of statuses excluding the current status *i*. The type of transition after the waiting time Δ*t* is then sampled randomly with probabilities proportional to the rates in *S*\*i*. We repeated these time-forward simulations for each species up to a pre-defined time of *t*_*max*_ = 100 years after present, producing 10,000 simulations for all 6 approaches outlined above: i) critE EX mode, ii) critE EX mode no GL, iii) critE EX mode no status change, iv) critE EX mode no GL and no status change, v) empirical EX mode, vi) empirical EX mode + PEX. For each simulation replicate, we repeated the drawing of a valid IUCN status for DD and NE species, thus incorporating this uncertainty in the simulations.

From the simulation output we extracted for each species a) the extinction times *t*_{*EX*}_ for those simulation replicates where *t*_*EX*_ < *t*_*max*_ or b) the waiting times of length *t*_*max*_ for those simulation replicates where the species did not go extinct within the specified time window. Next we used these extinction times and waiting times to estimate species-specific annual extinction rates averaged across the simulated time window.

For a given set of extinction times and waiting times simulated for species *i*, we applied a MCMC to obtain posterior samples of the extinction rate μ_*i*_ using the likelihood function of a death process (Silvestro et al. 2019):

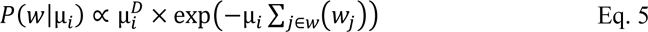

where *D* is the number of instances in which *w* ≤ *t*_*max*_, i.e. the number of simulation replicates in which the species went extinct within the considered time window.

We sampled estimates of μ_*i*_ for each species throughout the MCMC from the posterior distribution:

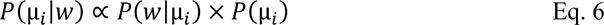

where *P*(*μ*_*i*_) is a uniform prior distribution (𝒰[0,1000]) set on the extinction rate. For each bird species we exported the mean and the 95% HPD interval of the posterior extinction rate estimates.

To compare the extinction rates between different approaches, we calculated the average extinction rate for each approach by running an MCMC with the death process likelihood function (Eq. 5), based on the simulated extinction dates of all bird species across 10,000 simulation replicates.

### Testing accuracy with synthetic data

In addition to the empirical bird data, we validated our approach on simulated data to determine the accuracy of the estimated transition rates and extinction rates.

To test the accuracy of the transition rates estimated from the IUCN history and the effect of the size of the chosen reference group on these estimates, we simulated status transitions data mimicking the empirical IUCN history data. We then simulated status changes over a time period of 20 years for reference groups of 100, 1,000, and 10,000 species. The starting status for each species was drawn randomly, based on the empirical frequencies of the current IUCN status distribution across all birds. To produce realistic transition rates to use for our simulations, we randomly drew these rates from a uniform range in log-space, ranging between the minimum and the maximum empirical rate estimated for birds. We drew 30 rates to reflect the 30 possible transition types between the six main IUCN statuses (LC, NT, VU, EN, CR, and DD). We then simulated the change of IUCN statuses through time in the same manner as described above for the future simulations for the empirical bird data, with the difference that no extinction events were being modeled. We then used the simulated IUCN history for all species to infer transition rates using MCMC as done for the empirical bird data.

To evaluate the accuracy of the species-specific extinction rates estimated with our approach, we simulated extinction times for 1,000 species under known extinction rates. The extinction rates (*μ*) that were used for these simulations were randomly drawn from a uniform range (in log-space) with a minimum and maximum value derived from the EX transition rates of the statuses LC and CR respectively, as modeled in this study with the outlined IUCN extinction probabilities approach. Based on the chosen number of simulation replicates, *N* extinction time replicates (*t*_*e*_) were drawn randomly from an exponential distribution with mean *μ*^−1^ for each species. This simulation was repeated for 100, 1,000, and 10,000 replicates, in order to test how many replicates are necessary for an accurate rate estimation.

## Results

### Transition rates

We counted a total of 919 status transitions between extant IUCN statuses in the IUCN history data of birds between the years 2001 and 2020 (Table 1). Additionally, we counted 6 transitions from CR to EX. This count increased to 20 when additionally modeling the PEX taxa as EX. The mean transition rates estimated from these counts, averaged across all 100 q-matrix replicates for all species, can be found in Table 2. With our transition rate estimation method, even transition types with zero-counts in the IUCN history are assigned a positive transition rate, although these rates will be very small. Differences between estimated rates can occur even if based on identical counts because of differences in the cumulative times spent in each category (see Eq. 2), as can be seen when for example comparing the transition rate from LC to EX with that from EN to EX, which differ by orders of magnitude (Table 2). A comparison between the estimated EX transition rates based on the two main approaches tested in this study (critE EX mode and empirical EX mode) show, that the empirical EX mode, which uses empirical extinction events, leads to lower average transition rates towards status EX for the threatened categories (Table 2).

The estimates based on our synthetic status transition data demonstrate that our approach accurately recovers the transition rates that were used to simulate the data, yet the precision of these estimates is strongly dependent on the size of the reference group (Fig. 3a). These results suggest that it is recommendable to choose reference groups of preferably more than 1,000 species, because stochastic fluctuations of status counts below that threshold preclude the estimation of transition rates with meaningful precision, particularly so for low rates.

**Figure 3:**
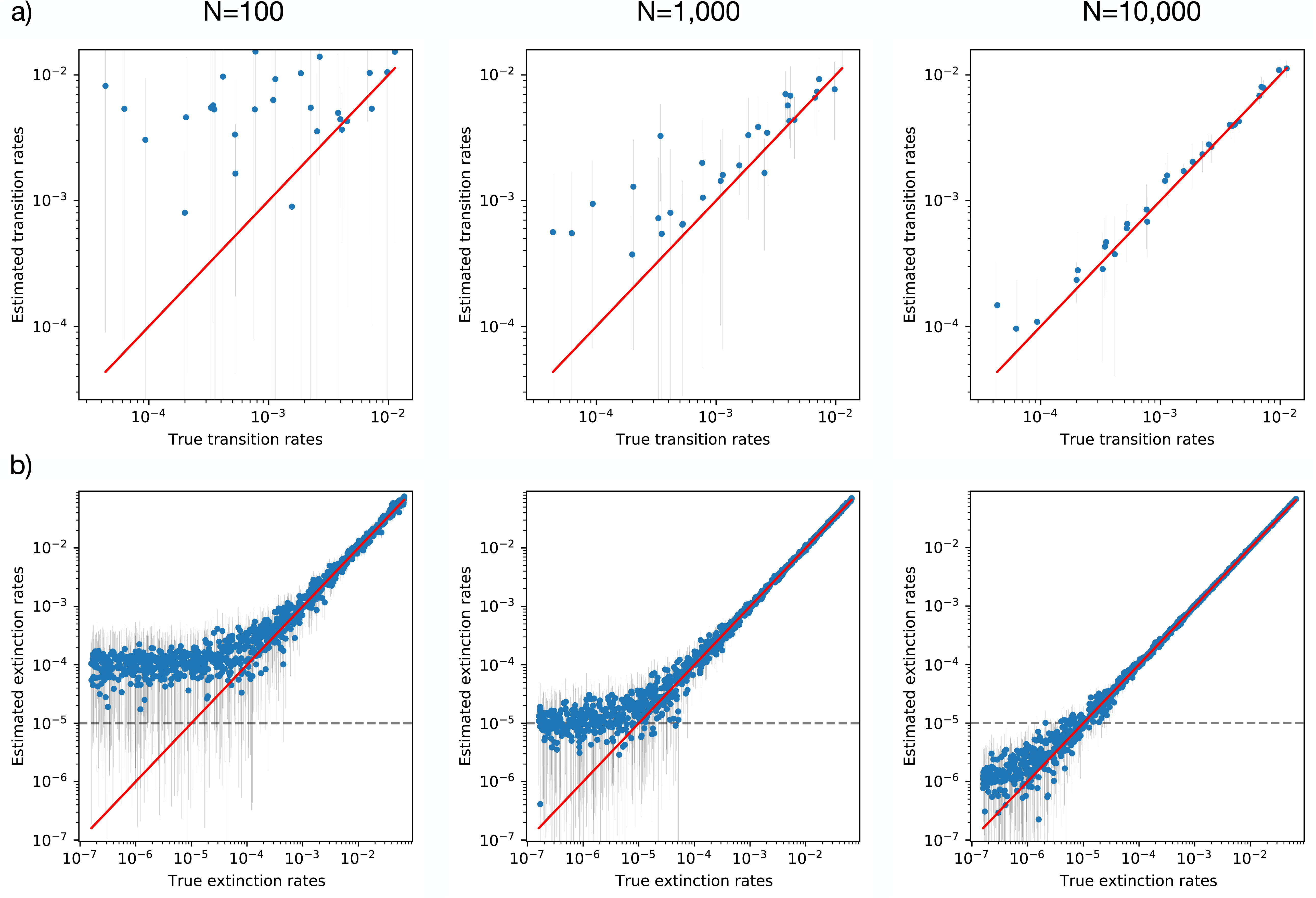
Accuracy of rate estimations improves with higher sample sizes. The scatter-plots show the status transition rates (a) and the extinction rates (b), estimated from synthetic data that was simulated in this study under known rates. We plotted the mean values (blue dots) and the 95% credible interval (grey vertical lines) of the rates sampled by MCMC (y-axis) against the true rates (x-axis) to evaluate the accuracy under different sample sizes (see plot titles). The sample size in case of the status transition rates (a) constitutes the number of species in the reference group, while sample size for extinction rates represents the number of future simulations for each species. Rate estimates close to the diagonal red line show high accuracy, while small error bars show high precision. Status transition rates estimated for reference groups of only 100 species show very low accuracy and therefore it is recommended to choose reference groups of at least 1,000 or more species. The dotted horizontal line in the extinction rate plots (b) shows the minimum empirical extinction rate estimate for the bird data (∼1 × 10^−5^). Extinction rates far below this line are therefore unlikely to occur in empirical data sets. Running at least 10,000 simulation replicates ensures accurate and precise extinction rate estimates.

### Future extinctions

Our future simulations for birds based on the critE EX mode approach resulted in 738 predicted species extinction within the next 100 years (95% confidence interval: 669-809 species, Fig. 2). In comparison the empirical EX mode approach resulted in 57 predicted extinctions within the same time frame (21-93), which increased to 127 (82-182) when accounting for PEX species. The species-specific extinction rates for all birds are available in the Supplementary Data.

The estimations of species-specific extinction rates from the simulated extinction times (Eq. 5) produces accurate rate estimates, yet it requires around 10,000 future simulation replicates to ensure this accuracy also for very low rates, such as species starting as LC (Fig. 3b). These species-specific rates differ significantly between the approaches tested in this study (Fig. 4). The empirical EX mode consistently leads to lower rate estimates than the critE EX mode, which is a direct result of the differences in EX transition rates in the q-matrix between these two approaches (Table 2).

**Figure 4:**
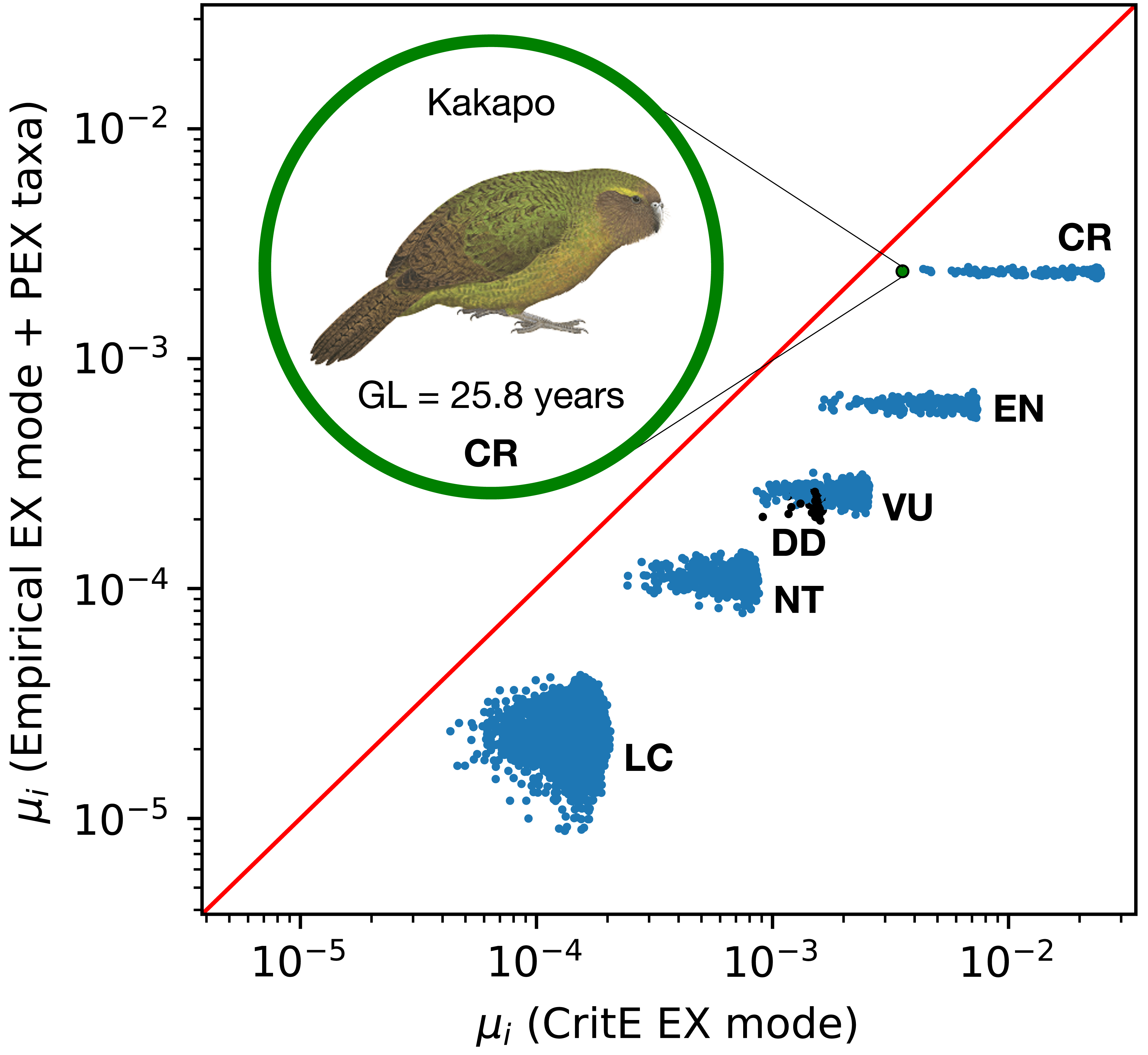
Species-specific extinction rates for the two ***iucn_sim*** approaches of modeling EX transition rates. The x-axis shows the estimated rates for the critE EX mode approach (all rates scaled in extinctions per species-years - ESY). The y-axis shows the estimated rates for the empirical EX mode approach including PEX taxa modeling. The rates estimated from the empirical EX mode approach are consistently lower than those from the critE EX mode approach. Species with the same IUCN status at present end up with similar rate estimates, forming visible clusters in the plot. However, there is some observed variation in the estimates between species of the same status, which is present in the estimates of both approaches (x and y-axis). This variation is partly caused by the stochasticity in our simulation approach. But particularly for the more threatened categories EN and CR we find additional variation among the critE EX mode rate estimates that is not present on the y-axis, causing the elongated shapes of these clusters as opposed to the round shapes of the less threatened status clusters. This is caused by differences in the GL values of individual species, leading to smaller extinction rate estimates for species with long generation times, as highlighted exemplarily for the CR Kakapo (*Strigops habroptila*), with one of the longest generation lengths in our dataset (25.8 years), which places on the very low rate end of the CR extinction rate cluster. In our approach DD species (black dots) are being modeled based on the observed DD transition rates in the IUCN history of the reference group, which in the case of birds results in extinction rate estimates similar to those of VU species. The illustration was provided by the Handbook of the birds of the world alive (Collar, N. et al. 2020).

The average rate estimated across all birds for the critE EX mode was 6.98 × 10^−4^ extinctions per species-years (ESY) (95% credible interval 6.97 − 6.98 × 10^−4^). This rate is to be understood as the average bird extinction rate expected over the next 100 years. The rate for the empirical EX mode was estimated to be significantly lower at 5.09 × 10^−5^ (5.08 − 5.11 × 10^−5^). When in addition modeling PEX taxa as extinct, the rate increased to 1.16 × 10^−4^ E/SY (1.15 − 1.16 × 10^−4^). These rate estimates fall within the same level of magnitude as previous estimates for birds, such as the 2.17 × 10^−4^ E/SY estimated by Monroe et al. (2019).

To further compare our results with those of Monroe et al. (2019), we additionally simulated extinctions and estimated extinction rates within a time window of 500 years, to match the time window addressed in their study, and we used the empirical EX mode setting to match the approach taken in their study. This resulted in an average rate estimate of 1.37 × 10^−4^ E/SY (1.369 − 1.370 × 10^−4^) for the empirical EX mode approach, which represents the average rate expected for the next 500 years under this model. These rate estimates are significantly lower than the 2.17 × 10^−4^ E/SY estimated by Monroe et al. (2019). Yet, the predicted number of extinctions under our approach ranged between 271 and 791, which largely overlaps with the 226 to 589 extinctions predicted by Monroe et al. (2019).

This discrepancy in rate estimates reflects a difference in how the rates are estimated and what they represent. Monroe et al. (2019) calculated their rate as the inverse of the average expected longevity (time until extinction) based on all birds. This corresponds to the average extinction rate of a process running until the extinction of all species. Our rate estimate, on the other hand, is based on simulated extinction events over the next 500 years and therefore reflects the average extinction rate within that time frame. Because in both approaches species threat statuses evolve according to an asymmetric transition matrix (Eq. 3, Table 2), the extinction process is not time-homogenous, as also noted by Monroe et al. (2019). Extinction rates increase through time as a consequence of the trend towards increasing frequencies of high threat statuses (Fig. 2). Consequently, rates averaged over a shorter time window (as the 500 years simulated in our case) are expected to be lower than rates averaged over the much longer time window reaching until the time of the expected extinction of all birds (Monroe et al. 2019).

Our rate estimates provide a representation of the extinction process specifically within the time window of interest and they change accordingly to the chosen simulation time. This is the reason why the rate estimates for the empirical EX mode approach, reported above, differ between the 100 year and the 500 years simulations (5.09 × 10^−5^, and 1.37 × 10^−4^ respectively).

To better compare the estimated rates between Monroe et al. (2019) and our study, we calculated the expected number of extinctions under each rate for the time interval of 500 years. Using the properties of a death process and assuming time-homogeneous rates within this interval we can compute the expected number of extinctions (D) based on a number *N* of initial species within a time interval *t* as:

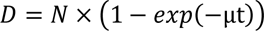

where the second term of the multiplication is the probability of surviving until time *t* given the extinction rate μ. Using this formula with *N* = 10,961 (number of extant bird species according to IUCN 2019-v2) and *t* = 500 years with our extinction rate (μ = 1.37 × 10^−4^) we obtain 726 expected number of extinctions, which is well within the range obtained from our simulations (271 - 791). In contrast, the 2.17 × 10^−4^ rate of Monroe et al. (2019) predicts 1,127 extinctions for the same time frame. This differs from their reported range of 226 to 589 expected extinctions, which was not estimated based on that reported rate, but derived from the expected longevities of all species based on an IUCN status transition q-matrix, similar to the one used in our study. Their rate estimate was calculated subsequently as the inverse of the average longevity across all birds and thus represents an overall rate averaged across the complete time frame until the extinction of all birds. Our rate estimate on the other hand is specific to the chosen simulation time window and describes more adequately the extinction process within that window. This demonstrates the utility of our **iucn_sim** program, which can be applied in future studies to predict extinction rates for specified time frames for any organism group or for individual species.

### Effect of modeling status change and GL data

Our empirical results show that accounting for GL data decreases the resulting extinction rate estimates (Fig. 5). As an example we highlight this effect for the Red-headed Vulture (*Sarcogyps calvus*), which is categorized as CR and has a relatively long generation length of 15 years (IUCN Red List 2019). The reduction of extinction probability when including GL is expected to be particularly strong for CR species with long GL times, since the immediate extinction probability applied in the simulations for EN and CR species decreases when incorporating the GL information, according to IUCN definition (critE EX mode). But the GL effect will also apply to LC species, as highlighted for the Turkey Vulture (*Cathartes aura*, GL = 9.9 years), where incorporating GL data leads to a small decrease in the extinction rate estimates, since occasionally these species will transition to the categories EN or CR in the future simulations, when allowing for future status changes (Fig. 5a). Overall, accounting for GL data leads to a decrease in the number of predicted extinctions across the whole target group (Fig. 6).

**Figure 5:**
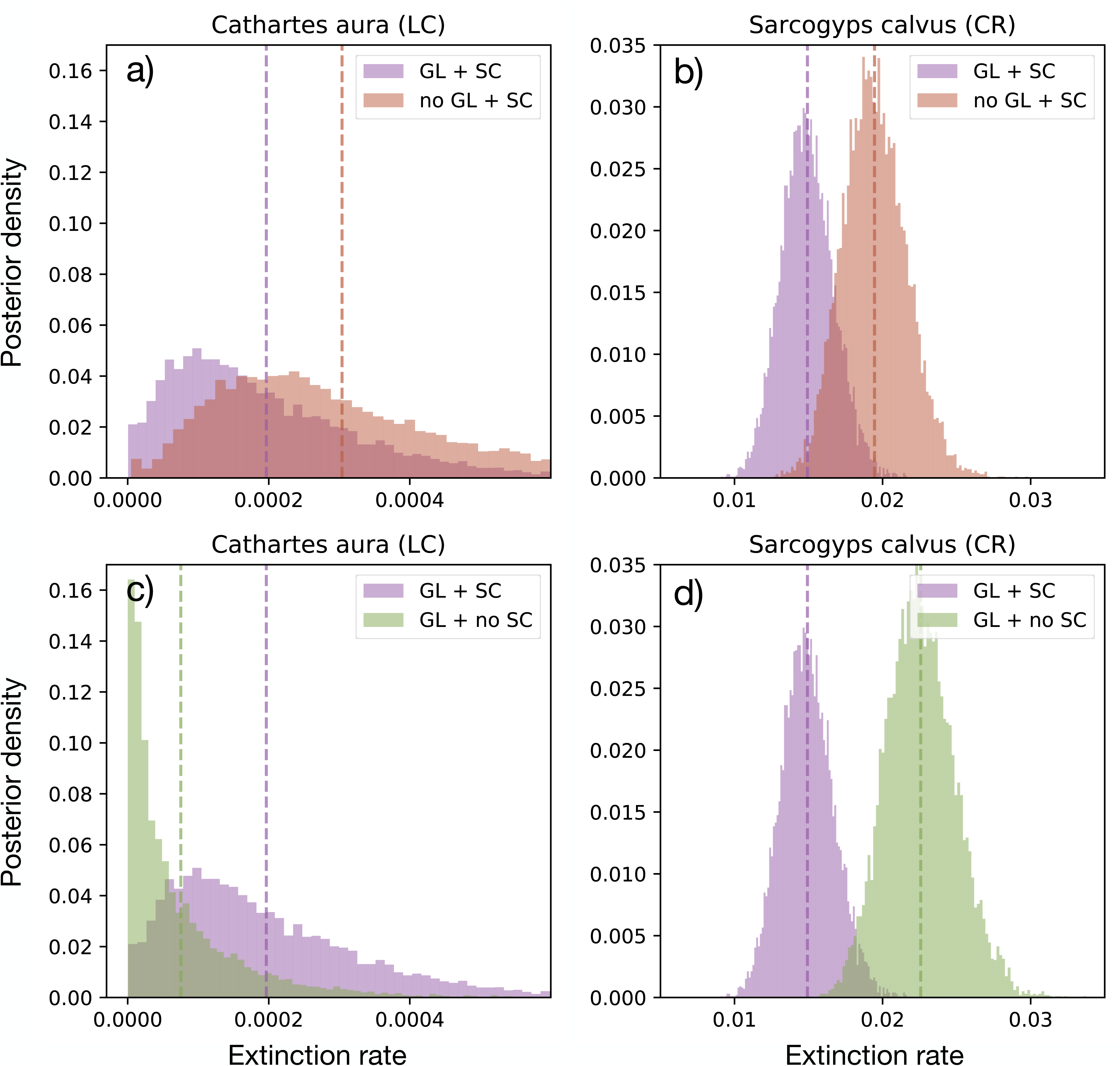
The effect of generation length (GL) and status-change (SC) on estimated extinction rates. The plots show histograms of the posterior density of extinction rates estimated with ***iucn_sim*** for two different species: the Turkey Vulture (Cathartes aura, GL = 9.9 years, LC), panels a) and c); and the Red-headed Vulture (Sarcogyps calvus, GL = 15 years, CR), panels b) and d). Upper panels show that the extinction rate estimates slightly decrease when including GL data into the simulations (purple) compared to ignoring GL data (red) for both LC and CR species. Bottom panels show that accounting for future changes of IUCN statuses slightly increases the extinction rate of LC species, but leads to a decrease for CR species (d). Note that the effect of future status changes on extinction rates depends on the estimated status transition rates and is therefore expected to change depending on the chosen reference group.

**Figure 6:**
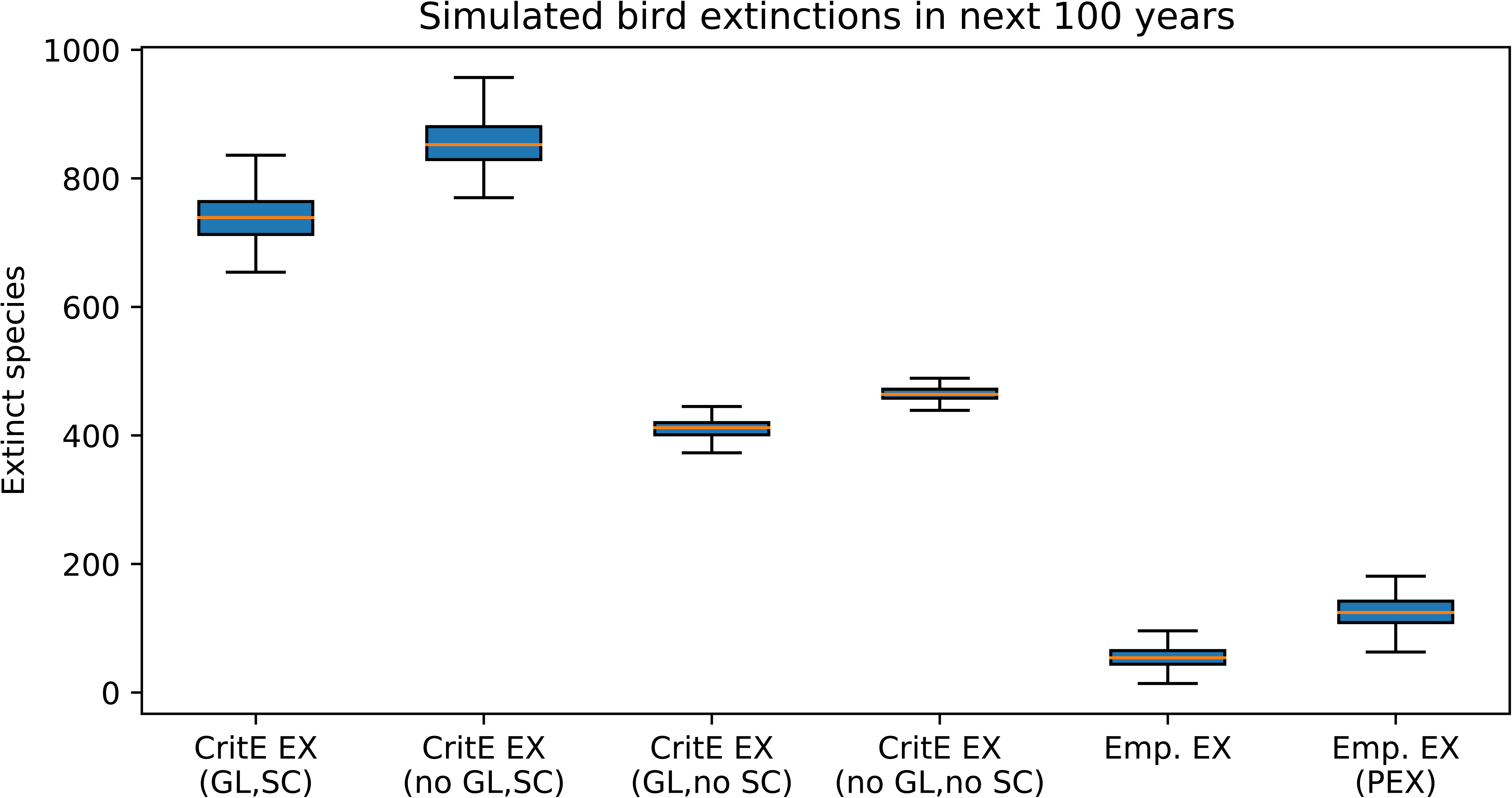
Number of predicted extinctions for birds in the next 100 years under different simulation scenarios across 100 simulation replicates. The blue boxes show the lower to upper quartile values of the predicted extinctions, with an orange line at the median. The whiskers show the full range of the predictions. Including generation length (GL) and conservation status changes (SC) into future simulations, leads to a significant increase in the number of predicted extinctions, compared to ignoring this information (compare first and fourth box plot column). The individual effect of adding GL information to the simulations is a decrease of the predicted extinctions (third box-plot column), while only modeling SC leads to very high numbers of predicted extinctions (second box-plot column). The last two columns show the range of predicted extinctions for the empirical EX mode approach, with (column 5) and without PEX taxa (column 6). The estimates for both empirical EX mode approaches are significantly lower than those for any of the variations of the critE EX mode approach (columns 1-4).

The effect of modeling future IUCN status changes can increase or decrease the estimated extinction rates of a species, depending on its current status and on the transition rates between statuses. Therefore, this effect is expected to be dependent on the chosen reference group. However, for LC species this generally appears to lead to an increase in the estimated extinction rates (Fig. 5c), which is likely because these species can only change to a more threatened status (LC being the least threatened status). Similarly, for CR species, the effect of modeling future status changes generally leads to a decrease in extinction rates (Fig. 5d), since species can only switch to less threatened categories in the future (CR being the most threatened status). Overall, modeling future status changes leads to a sharp increase in the number of predicted extinctions across the whole target group (Fig. 6), compared to the scenario with no future status changes.

## Discussion

### Utility of the iucn_sim program

With our open-source program ***iucn_sim*** that accompanies this study, we are presenting improved versions of the two main approaches of previous studies for modeling future biodiversity losses based on IUCN status assessments (Fig. 1). Through this, we hope to facilitate future studies to apply these workflows for generating future diversity predictions and for estimating extinction rates for whole groups or individual species. The program is easy to use and to simulate future extinctions it requires only a list of target species names, or even just the name of the taxonomic group, as it automatically retrieves all available IUCN information. Moreover, ***iucn_sim*** also allows for additional data input for more specific estimates, such as GL data, that the user can choose to provide.

One of the main outputs of the program is the predicted number of future species extinctions for a given group of species, as well as the future changes of the IUCN status distribution within the group (Fig. 2). The program also calculates the probabilities of each species being extinct by a user-defined date. Finally, the program estimates the extinction rates based on the simulated extinction dates separately for each species (Fig. 4). These species-specific extinction rates can be of interest for downstream analyses where species are required to be modeled individually based on biological or geographic data, and where the extinction dynamics of specific species or groupings of species are of interest (Davis et al. 2018, Cooke et al. 2019, Pimiento et al. 2020).

We note that the actual extinction rates of a given species or group are expected to vary over time as a function of changes in the IUCN status, while the extinction rates inferred by ***iucn_sim*** are a time-averaged proxy of this process. Particularly during the current human-induced wave of extinctions, extinction rates are expected to vary within relatively short time frames of at least 100s of years (Ceballos et al. 2015). Therefore, our approach presented here may not be suitable for estimating extinction rates based on simulations that span across several hundred years or more.

Our method further allows for modeling DD species for which IUCN statuses are imputed based on historical transition rates that reflect how frequently DD species change to other statuses. Similarly, a new status for NE species is modeled based on the current status distribution of the reference group. As a status is imputed for DD and NE species at each simulation replicate, our method incorporates the full uncertainty concerning their true status. The ***iucn_sim*** program flags and prints to the screen the names of those species that cannot be found in the IUCN taxonomy and produces a warning for the user to revise the taxonomy. If these cases cannot be fixed by the user, they will be treated as NE. This approach enables future simulations even for groups where it is difficult to match the taxonomy with that of IUCN, yet we recommend thoroughly revising the taxonomies to minimize the number of taxonomic mismatches. Never the less, species unknown to IUCN, which are modeled in this manner, are not expected to bias the overall future biodiversity predictions (under the assumption that these taxa constitute a random sample of the target species group), due to their status being repeatedly resampled based on the empirical status distribution of the reference group. While these species are not expected to affect the overall predictions for the target group, the resulting species-specific extinction rates for these taxa on the other hand may be misrepresentative. To address this issue, the user can manually change the status of NE species to a status they deem more representative for the species, by altering the status in the *species_data.txt* text file produced by ***iucn_sim***.

Our ***iucn_sim*** program further allows the simulation of different future conservation scenarios, through simple q-matrix modifications. For example, one can simulate an increase of conservation efforts by providing a specific conservation factor. This factor is then applied to all transition rates in the q-matrix, leading to an improvement in conservation status for each species. Similarly, one can simulate increased threats by providing a threat factor, which is then applied to all threat-increasing transition rates in the q-matrix. These factors can also be set to 0 to simulate scenarios that do not allow for future improvements or increased threats. This flexibility of ***iucn_sim*** makes it easy to simulate and compare different future scenarios and their expected effect on biodiversity.

### Comparing approaches to simulate future extinctions

The critE EX mode and empirical EX mode approaches that were applied in this study represent different ways of modeling the EX transition rates, which are the rates at which species transition towards extinction in our future simulations. These are not to be confused with the species-specific extinction rates, which are instead estimated from the simulated extinction times and describe the extinction risk of individual species.

The critE EX mode makes use of extinction probabilities that are defined by the IUCN as one of several criteria for species assessments of threatened species. Although widely used in the scientific literature for modeling species’ extinction risks (Veron et al. 2016, Davis et al. 2018, Cooke et al. 2019, Oliveira et al. 2019), these probabilities are not originally intended for this purpose and per definition only apply to the subset of threatened species that was assessed under criterion E (Akçakaya et al. 2006). The simulated extinctions resulting from this approach are alarmingly high and the estimated extinction rates are in most cases more than an order of magnitude higher than those estimated with the empirical EX mode approach, even when accounting for PEX taxa in the latter approach.

The empirical EX mode, on the other hand, will likely lead to an underestimation of the true extinction rates, because it is directly dependent on the number of observed transitions from extant categories to EW or EX in the IUCN history, and these documented numbers are likely a significant underestimate (IUCN 2020). This underestimation bias is due to rather strict requirements to classify species as EW or EX. In 2020, IUCN therefore released a list of species that are possibly extinct (PEX species), but do not qualify as EX according to the IUCN guidelines. Making use of this information (which is available in ***iucn_sim***) and modeling these taxa as extinct, usually leads to more observed status transitions towards EX within the last 20 years of IUCN history and therefore leads to higher EX transition rate estimates that are expected to better reflect the true extinction risk within the studied group. However, this approach is expected to be sensitive towards small reference groups with very few or no observed extinctions, which will lead to high uncertainties in the estimated rates. Given these significant differences in predicted future estimates between the two approaches, it is important to consider that these approaches are based on different assumptions and while both can theoretically be applied for any organism group, their utility varies depending on the group and purpose of the future simulations.

If the primary aim is to conservatively model future biodiversity losses for a given group of species, and if this group can be meaningfully represented by a reference group that is a) well represented in the IUCN Red List (i.e. many assessed species), b) has a high species diversity, and c) has several recorded extinctions throughout the last 20 years, then the empirical EX mode including PEX taxa may be the most suitable choice, as in this case EX transition rates can be meaningfully modeled for the specific reference group, rather than being based on general pre-defined extinction probabilities.

If, on the other hand, the primary aim is to produce species-specific extinction rates for downstream analyses, and GL data is available or can be modeled for the group of species, then the critE EX mode approach may be the more appropriate choice, as it leads to a larger variation of rate estimates. This variation is expected to reflect differences in how threatened with extinction species of the same category are, based on their differences in GL. In the empirical EX mode approach on the other hand all species belonging to a given category are modeled equally, only leading to small stochastic differences between the rates of species belonging to the same status.

### Choice of reference group

Essential to both approaches discussed above is the choice of the reference group, because the precision and accuracy of the estimated transition rates depends on the number of species in the reference group (Fig. 3). There are two main considerations to make when choosing a reference group: 1) Is the chosen group expected to reflect the trends of status change for the target species that are being simulated? and 2) Does the reference group contain a sufficient number of species so that stochastic effects do not overrule the actual trends for that group?

These two objectives can conflict, as illustrated by the example of simulating future extinctions for vultures. In that case, using all birds (class Aves) as reference group (∼ 11,000 species) provides a large enough group of sufficient size for accurate transition rate estimations. However, given the notable recent worsening of almost all vulture species’ conservation status (e.g. Green et al. 2007), the trends observed over all birds may not be representative of this group.

The species in the reference group do not necessarily have to form a monophyletic clade, although phylogenetically related taxa are likely to provide a suitable reference group if there is any phylogenetic signal in extinction risk. More importantly, a suitable reference group consists of species that are expected to share the same extinction threats as the group of target species for which to simulate future extinctions, so that representative status transition rates can be inferred. A reference group could include species that share a similar ecology and are similarly affected by habitat losses or pollution, or it could include species from the same biogeographic area as the target species if they are expected to share common threats, such as is the case for many island faunas. For these reasons, the reference group should also always contain all of the target species, although this is not an analytical requirement.

## Conclusions

In this study, we demonstrated that modeling future changes in IUCN conservation status and incorporating generation length data has a substantial effect on future extinction predictions. In addition, we encountered significant differences in extinction rate predictions when comparing different approaches of modeling extinction risks. This shows that the results of future projections are strongly dependent on the selected method and the selected reference group.

The aim of this study was to make the simulation of future extinctions under different approaches accessible for future projects. Further, this study can provide a starting point for researchers to decide which approach to choose for their specific target group and research objective.

Our software ***iucn_sim*** is designed for ease of use and contains many options for adjusting the simulation approach for different types of projects. The source code on GitHub is open for contributions and feedback from users, which hopefully will lead to the incorporation of further improvements for predicting future extinctions. Future additions to the program could for example include more specific future modeling of species based on similarities in biological traits, geographic location, or niche space.

### Data availability statement

All source code, input files used in this study, and the output produced by ***iucn_sim*** are available on the project’s GitHub repository at https://github.com/tobiashofmann88/iucn_extinction_simulator. The estimated extinction rates for all bird species, a Supplementary Code Sample describing the ***iucn_sim*** workflow, and Appendix 1, can be directly downloaded from the Supplementary Material accompanying this study.

## Supporting information

Code Sample 1

## References

Akçakaya, H. R. et al. 2006. Use and misuse of the IUCN Red List Criteria in projecting climate change impacts on biodiversity. - Global Change Biology 12: 2037–2043.

Barnosky, A. D. et al. 2011. Has the Earth’s sixth mass extinction already arrived? - Nature 471: 51–57.

Bird, J. P. et al. 2020. Generation lengths of the world’s birds and their implications for extinction risk. - Conservation Biology in press.

BirdLife International 2019. BirdLife Data Zone. Available at http://datazone.birdlife.org/home.

Butchart, S. H. M. et al. 2007. Improvements to the Red List Index (D Lusseau, Ed.). - PLoS ONE 2: e140.

Ceballos, G. et al. 2015. Accelerated modern human – induced species losses: entering the sixth mass extinction. - Science Advances 1: 1–5.

Chamberlain, S. 2017. Rredlist: “IUCN” Red List Client.

Collar, N. et al. 2020. Kakapo (Strigops habroptila). - In: Handbook of the Birds of the World Alive. Lynx Edicions, Barcelona, in press.

Cooke, R. S. C. et al. 2018. Improving generation length estimates for the IUCN Red List (L Bosso, Ed.). - PLOS ONE 13: e0191770.

Cooke, R. S. C. et al. 2019. Projected losses of global mammal and bird ecological strategies. - Nature Communications 10: 2279.

Davis, M. et al. 2018. Mammal diversity will take millions of years to recover from the current biodiversity crisis. - Proceedings of the National Academy of Sciences of the United States of America 115: 11262–11267.

Díaz, S. et al. 2019. Summary for policymakers of the global assessment report on biodiversity and ecosystem services. - Intergovernmental Science-Policy Platform on Biodiversity and Ecosystem Services, IPBES.

Frankham, R. and Brook, B. W. 2004. The importance of time scale in conservation biology and ecology. - Annales Zoologici Fennici 41: 459–463.

Green, R. E. et al. 2007. Rate of Decline of the Oriental White-Backed Vulture Population in India Estimated from a Survey of Diclofenac Residues in Carcasses of Ungulates (D Carter, Ed.). - PLoS ONE 2: e686.

IUCN 2020. Summary Statistics - Table 9: Possibly Extinct species.

IUCN Red List 2019. Red List of threatened species v2019-2. Available at http://www.iucnredlist.org. - Version 2019-2

IUCN Species Survival Commission 2001. IUCN Red List Categories and Criteria: Version 3.1. Available at https://portals.iucn.org/library/sites/library/files/documents/RL-2001-001.pdf.

IUCN Standards and Petitions Committee 2019. Guidelines for Using the IUCN Red List Categories and Criteria. Version 14. Available at http://www.iucnredlist.org/documents/RedListGuidelines.pdf.

Kindvall, O. and Gärdenfors, U. 2003. Temporal Extrapolation of PVA Results in Relation to the IUCN Red List Criterion E. - Conservation Biology 17: 316–321.

Monroe, M. J. et al. 2019. The dynamics underlying avian extinction trajectories forecast a wave of extinctions. - Biology Letters 15: 20190633.

Mooers, A. Ø. et al. 2008. Converting endangered species categories to probabilities of extinction for phylogenetic conservation prioritization. - PLoS ONE 3: 1–5.

Oliveira, B. F. et al. 2019. Decoupled erosion of amphibians’ phylogenetic and functional diversity due to extinction (A Ordonez, Ed.). - Global Ecology and Biogeography: geb.13031.

Pacifici, M. et al. 2013. Generation length for mammals. - Nature Conservation 5: 89–94.

Pimiento, C. et al. 2020. Functional diversity of marine megafauna in the Anthropocene. - Science Advances 6: eaay7650.

Redding, D. W. and Mooers, A. Ø. 2006. Incorporating Evolutionary Measures into Conservation Prioritization. - Conservation Biology 20: 1670–1678.

Ricciardi, A. and Rasmussen, J. B. 1999. Extinction Rates of North American Freshwater Fauna. - Conservation Biology 13: 1220–1222.

Rondinini, C. et al. 2014. Update or Outdate: Long-Term Viability of the IUCN Red List. - Conservation Letters 7: 126–130.

Silvestro, D. et al. 2019. Improved estimation of macroevolutionary rates from fossil data using a Bayesian framework. - Paleobiology 45: 546–570.

Swinnerton, K. J. 2001. The ecology and conservation of the pink pigeon Columba mayeri in Mauritius.

Veron, S. et al. 2016. Integrating data-deficient species in analyses of evolutionary history loss. - Ecology and Evolution 6: 8502–8514.

