## Appendix 1 for "*iucn_sim*: A new program to simulate future extinctions based on IUCN threat status"

**Appendix 1: Phylogenetic imputation of generation length data**

This represents an example of how we estimated generation length data (GL) for all Aves species that did not have such information available. The data produced from this workflow was not included in the study, but instead we downloaded peer-reviewed estimates of GL from (Bird et al. 2020).

**The hypothetical workflow:**

We gathered GL estimates for all bird species to apply these for modeling species-specific extinction probabilities, following the IUCN v2019-2 taxonomy of extant bird species (IUCN Red List 2019). Generation length data for the majority of these species was provided by BirdLife International (http://www.birdlife.org). For all remaining species we modeled GL estimates using multivariate phylogenetic imputation under the assumption that GL has a phylogenetic correlation and is also correlated with body mass. Body mass data was downloaded from Cooke et al. (2019), which is based on data from the databases EltonTraits (Wilman et al. 2014) and the Amniote Life History Database (Myhrvold et al. 2015). For the phylogenetic imputation, we downloaded 1,000 samples of the posterior species trees distribution produced by (Jetz et al. 2012), based on the Ericson backbone ("EricsonStage2_0001_1000.zip"). A fraction of 90% of bird species names listed in IUCN v2019-2 were also present in the phylogenies. After manual taxonomic revision, we matched 96% of all IUCN bird species with the tips in the phylogenies.

To estimate GL values for all species lacking such data, we ran a phylogenetic imputation, using the R-package ***rphylopars*** (Goolsby et al. 2017). To determine the best model, we calculated the Akaike information criterion score for all available models in ***rphylopars*** (Fig. 1) and chose the early burst ("EB") as the best fitting model. In order to incorporate the uncertainty of the phylogenetic estimates, we ran separate imputations for 100 randomly selected trees from the downloaded species tree distribution. We exported the 100 resulting mean values of the GL estimates for each species.

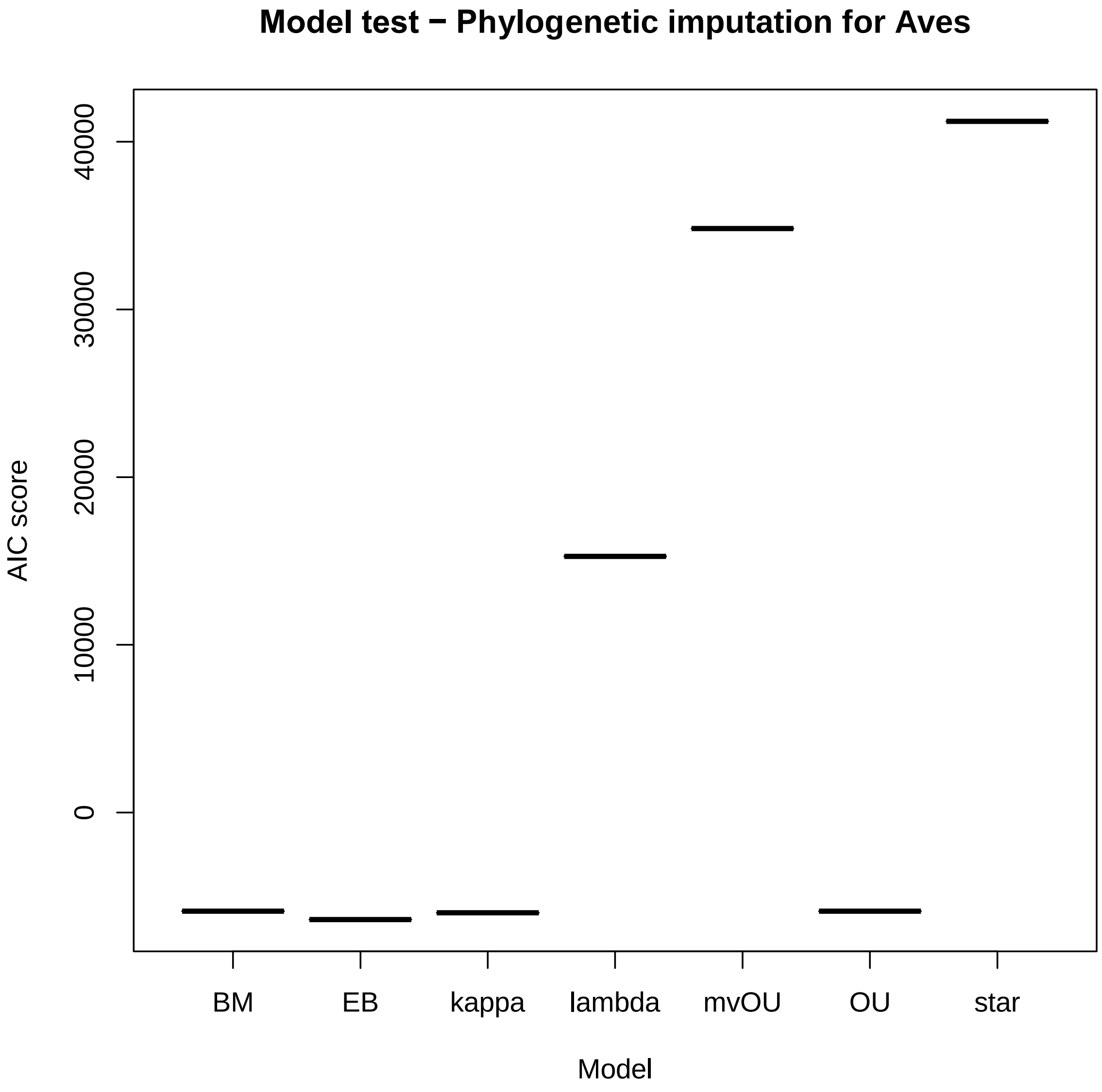


**Figure 1**: Results of model test in rphylopars. The plot shows the AIC scores of all tested models for the multivariate phylogenetic imputation of GL in birds. Based on these results we selected 'EB' as the best model.

For all remaining species that were not present in the phylogeny we used mean GL across their genus (congeneric mean GL), calculated separately for each of the 100 GL data replicates. This resulted in our final dataset containing GL estimates for all bird species listed by IUCN v2019-2.

**References**


Bird, J. P. et al. 2020. Generation lengths of the world's birds and their implications for extinction risk. - Conservation Biology in press.

Cooke, R. S. C. et al. 2019. Projected losses of global mammal and bird ecological strategies. - Nature Communications 10: 2279.

Goolsby, E. W. et al. 2017. Rphylopars: fast multivariate phylogenetic comparative methods for missing data and within-species variation. - Methods in Ecology and Evolution 8: 22–27.

IUCN Red List 2019. Red List of threatened species v2019-2. Available at http://www.iucnredlist.org. - Version 2019-2

Jetz, W. et al. 2012. The global diversity of birds in space and time. - Nature 491: 444–448.

Myhrvold, N. P. et al. 2015. An amniote life-history database to perform comparative analyses with birds, mammals, and reptiles. - Ecology 96: 3109–000.

Wilman, H. et al. 2014. EltonTraits 1.0: Species-level foraging attributes of the world's birds and mammals. - Ecology 95: 2027–2027.
