## Supplementary material for "*iucn_sim*: A new program to simulate future extinctions based on IUCN threat status": Code Sample 1

```bash
# Download IUCN history data, current status list, and list of PEX taxa
iucn_sim get_iucn_data \
    --reference_group aves \
    --reference_rank class \
    --target_species_list data/precompiled/gl_data/aves_gl.txt \
    --outdir data/iucn_sim_output/aves/iucn_data \
    --iucn_key <IUCN-key>

# Produce q-matrices, critE EX mode
iucn_sim transition_rates \
    --species_data data/iucn_sim_output/aves/iucn_data/species_data.txt \
    --iucn_history data/iucn_sim_output/aves/iucn_data/AVES_iucn_history.txt \
    --extinction_probs_mode 0
    --rate_samples 100
    --seed 1234
    --outdir data/iucn_sim_output/aves/transition_rates_0

# Produce q-matrices, empirical EX mode
iucn_sim transition_rates \
    --species_data data/iucn_sim_output/aves/iucn_data/species_data.txt \
    --iucn_history data/iucn_sim_output/aves/iucn_data/AVES_iucn_history.txt \
    --extinction_probs_mode 1
    --rate_samples 100
    --seed 1234
    --outdir data/iucn_sim_output/aves/transition_rates_1

# Produce q-matrices, empirical EX mode + PEX
iucn_sim transition_rates \
    --species_data data/iucn_sim_output/aves/iucn_data/species_data.txt \
    --iucn_history data/iucn_sim_output/aves/iucn_data/AVES_iucn_history.txt \
    --extinction_probs_mode 1
    --possibly_extinct_list data/iucn_sim_output/aves/iucn_data/
        possibly_extinct_reference_taxa.txt \
    --rate_samples 100
    --seed 1234
    --outdir data/iucn_sim_output/aves/transition_rates_1_pex

# Run future simulations and estimate extinction rates (replace XXX)
iucn_sim run_sim \
    --input_data data/iucn_sim_output/aves/transition_rates_XXX/
        simulation_input_data.pkl
    --outdir data/iucn_sim_output/aves/future_sim_XXX
    --n_years 100
    --n_sim 10000
    --extinction_rates 1
    --seed 1234
```

Code sample 1: The `iucn_sim` commands corresponding to the methods described in the main text. These commands can be executed in the bash command line on any operating system, after installing `iucn_sim` with the conda reference manager. For installation instructions and a more comprehensive tutorial, visit the GitHub page of this project (`https://github.com/tobiashofmann88/iucn_extinction_simulator`). The only required input file in this case is the `aves_gl.txt` file, which can be found together with all output files of this workflow in the projects GitHub repository under the paths specified in the respective commands. The input files for all other functions are files produced by `iucn_sim`. For running the `get_iucn_data` function, a IUCN API key is required, which can be requested for free from IUCN (`https://apiv3.iucnredlist.org/api/v3/token`).
